# *Hoslundia opposita* and other Nigerian plants inhibit sickle hemoglobin polymerization and prevent erythrocyte sickling: a virtual screening-guided identification of bioactive plants

**DOI:** 10.1101/2021.07.25.453693

**Authors:** Eric O. Olori, Olujide O. Olubiyi, Chinedum P. Babalola

## Abstract

In sickle cell disease, a hereditary hemoglobinopathy, clinically observed disease presentations are the endpoint of a point mutation involving the substitution of glutamic acid with valine at the position 6 of the beta globin chain. With about 4.4 million people globally being affected, and another 43 million people bearing the sickle cell trait, several research efforts have been made to discover new and affordable treatment and possibly cure for the disease. Africa is endowed with a large flora population and traditional healers and citizens have over time depended on the use of herbs in folkloric medicine for different ailments including sickle cell disease (SCD). Such native knowledge has often formed the basis for different research exploration into the anti-sickling activities of selected African plants. These plants that have been so far investigated for their anti-sickling properties represent about 0.05 % of the 45,000 plant species enriching the flora landscape in Sub-Saharan Africa. Some of these have yielded potent anti-sickling profiles. In the current work we seek to achieve a more extensive search of the African plant diversity with anti-sickling properties: for this we have adopted a hybrid computational-cum-experimental protocol that employed computer-aided drug design (CADD) means for identifying plants with at least one constituent capable of interacting with the sickle hemoglobin, followed by extractive procedures and anti-sickling experiments for validating the predictions. Over two thousand (or 2,000) African natural products, representing over 200 plant species, were first virtually screened against the crystal structure of the dimerized human sickle hemoglobin. The natural products with the best computed sickle hemoglobin interaction energetics were found to belong to five plant species including *Catharanthus roseus, Rauvolfia vomitoria, Hoslundia opposita, Lantana camara and Euphorbia hirta*. The leaves of these plants were each collected subsequently and subjected to standard processing and extraction procedures. Using both HbSS polymerization inhibition and sickling reversal tests significant reductions in polymerization of erythrocyte hemolysate of the HbSS genotype were observed with the methanolic extracts of the plants, as well as sickling reversal levels of up to 68.50 % (*H. opposita*) was observed.

## Introduction

Sickle cell disease (SCD) is a blood disorder affecting the red blood cells and is inherited (NHLBI, 2015). The disease was reported to be majorly prevalent in resource limited countries and about 80% of SCD cases occur in sub-Saharan Africa (Rees, 2010). Globally about 4.4 million people have SCD while 43 million bear the sickle cell trait (GBD, 2015) In 2010, the birth of children with sickle cell anemia (SCA) accounted for 2.4% the world’s most severe cases of the disease (Piel FB et al, 2013a). However, worrying estimates indicate that the number of newborns with SCA will increase from approximately 305,000 in 2010 to 404,000 in 2050 (Piel FB et al, 2013b & Piel FB et al, 2017). About 300,000 babies are born worldwide with sickle cell disease each year, spread across Africa and India (Piel et al, 2015). In 2010, 90,000 births with sickle cell disease were estimated in Nigeria and 40,000 in the Democratic Republic of Congo (Piel et al, 2013). Another estimated 40,000 children affected with sickle cell disease are born in India each year, 10,000 in the Americas, 10,000 in the Eastern Mediterranean, and 2000 in Europe (Piel FB et al, 2013). Newborn screening programs in the United States, Jamaica, and Europe have documented the utility of early identification of SCA, with a marked reduction in morbidity and mortality, especially in the first 5 years of life (Rogers et al. 1978; Vichinsky et al. 1988; Almeida et al. 2001; Bardakdjian-Michau et al). Sickle cell anaemia resulting from SCD is characterized by abnormality in the oxygen carrying haemoglobin (haemoglobin S, HbS) which at the molecular level results from a point mutation replacing a charged and hydrophilic glutamic acid with hydrophobic valine at the position 6 of the beta globin polypeptide chain (Pauling L et al, 1949 & Ingram VM, 1957). Under deoxygenation conditions (low oxygen tension), the mutant HbS polymerizes inside the red blood cells (RBCs) into a stiff gel and further into fibers leading to a decrease in red blood cell (RBC) deformability. HbS polymerization within erythrocytes provokes an alteration in the functional shape of red cells from the normal spherical form into distorted shapes some of which resemble a sickle (Bisnu et al, 2013 & Herrick, 1910). With the HbS molecule occupying a central position in disease development, it makes sense that efforts to evolve new leads for SCD management should, as a viable option, consider the inhibition of HbS polymerization both by small molecules (Ismaila O. Nurain, 2017) and larger molecular weight peptide-based inhibitors (Olubiyi et al, 2019). For instance, the recently FDA-approved voxelotor is a covalent modifier of HbS molecule and acts as an allosteric modulator stabilizing the hemoglobin molecule in its non-aggregating R-conformer (Vichinsky et al, 2019).

Normal hemoglobin has a quaternary structure existing as a tetrameric protein unit with two alpha- and two beta-chains (Robert J et al, 2014) with each of the four chains covalently linked to a heme molecule. There are 141 and 146 amino acids in the α and β chains of hemoglobin, respectively (Robert J et al, 2014). While interactions between multiple hemoglobin tetrameric units can in principle result in polymerization, such resulting aggregates are however unstable under normal physiological conditions. In homozygous SS patients on the other hand, the beta-chains contain valine at the sixth position which is able to form hydrophobic contacts with similarly uncharged hydrophobic residues like Phe85 and Leu88 of the adjacent beta globin chain. These contacts constituting the lateral HbS contacts are secondarily stabilized by other contacts involving segments of the hemoglobin molecule that are remote from the valine 6 mutation. Taken together, these contacts constitute the basis for HbS polymerization and by extension SCD symptom complex development. Within the same beta-2 globin chain of an HbS molecule there is glutamic acid at position 121 which interacts with Gly16 of the beta-1 globin chain of another HbS molecule. And, between these two Hb molecules, His20 from alpha-2 globin chain interacts with Glu6 of beta-1 of the third hemoglobin molecule as part of the axial contacts. Apart from beta-2 globin’s Val6 interacting with Phe85 and Leu88 of the adjacent beta globin chain, Asp73 and Thr4 from the two beta globin chains are also involved in contacts (Marengo-Rowe, 2006 & Levasseur DN et al, 2004) Glu121 from beta-1 chain and Pro114 of alpha-2 and His116 of beta-2 globin chain also form interaction (Ferrone, F. A et al, 1985 & Ivanova, M, 2002). Within the lateral contact, the hydrophobic Val6’s side chain fits well into a hydrophobic pocket formed by β-Leu88 leucine and β-Phe85 residues on an adjacent beta globin chain (Ivanova M et al, 2000).

Sickle cell disease is embodied by serious hemorheological irregularities which play a role in the pathophysiology of various severe and chronic complications. Normal RBC are pliable, able to squeeze through tight capillary junctions and can live for 90 to 120 days. During a typical 120 days lifespan of a RBC, it circulates through arteries, veins and small capillaries traveling –in total– a distance of 500 km (Lasch et al., 2000). Because sickle RBCs have stiffened texture they cannot as readily change their shape; the stress of squeezing through tight vascular junctions contributes to their tendency to hemolyze and as a result sickle RBCs last only 10 to 20 days, hence the associated anemia. Common signs and symptoms of the disease include painful swelling (hands and feet; vaso-occlusive crisis), fatigue from anemia (aplastic crisis), yellowing of the skin and white of the eyes, infections (splenic sequestration crisis), fever, chest pain and difficulty in breathing (acute chest syndrome), hemolytic crisis (Serjent GR, 2001 & Kar BC, 1991). While bone marrow cell transplant is the only available curative treatment of SCD (NHLBI, 2002), the following treatments are also available in the management of SCD: vaccination, antibiotics (for the infection component), high fluid intake, folic acid supplementation analgesic (e.g. Ibuprofen), hydroxyurea, blood transfusion and herbal preparations. Hydroxyurea has been proven to decrease the number of SCD crisis (American Society, 2016). Mechanistically it stimulates the production of fetal hemoglobin (HbF) and which inhibits HbS aggregation and has also been found to reduce white blood cells component of the inflammation crisis (Lanzkron S et al, 2008). The health-care cost of the management of SCD patients is disproportionately high compared to the number of people afflicted by the disease and considering that most affected individuals are below the poverty line and unable to afford the high-cost of treatment. This in particular creates the urgent need for the unraveling of potentially cheap natural products and plants-based treatment options. (Bisnu et al, 2013). One of such efforts produced Niprisan™, a herbal preparation developed at the Nigerian Institute for Pharmaceutical Research and Development (NIPRD) and indicated for the treatment of SCD. Niprisan was formulated from the ethanolic extract of *Piper guineense* seeds, *Pterocarpus osum* stem, *Eugenia caryophylius* fruit and *Sorghum bicolor* leaves. About 45,000 plant species are found in Sub-Saharan Africa (Klopper, R. R et al, 2007) and only about 0.05% has been researched and found to possess desirable antisickling properties. According to Sunday J. Ameh, discovery of more effective antisickling plants from the largely untapped African flora wealth will require the use of more innovative strategies (Ameh et al, 2012). It is certainly not far-fetched to expect that many more plants with more potent antisickling components exist in the African floral population that are yet undiscovered and the challenge remains how to efficiently identify these in eco-compatible ways. In this work we propose a protocol relying on the structural understanding of HbS aggregation for targeted screening of natural products from African plants against a virtual model of the HbS macromolecule. Our design strategy is based on the double nucleation aggregation mechanism of the HbS molecule. We believe that natural products capable of forming a precise fit into Val6 binding cavity on the lateral aggregation surface of the HbS molecule will be able to delay the formation of a critical HbS nucleus (homogenous nucleation), which in turn is capable of preventing template-driven nucleation altogether and thus preventing aggregation. Using this protocol we identified Nigerian plants containing the natural products shown in our computational drug design work; these were collected and subjected to in vitro antisickling and polymerization reversal assays. To the best of the knowledge of the authors, this work represents the first computer-guided selection of plants for screening of plants for antisickling activities.

Many researches have been conducted by various researchers in recent times to find new drugs from natural products for the treatment of SCD using traditional screening methods of drug discovery. These screening methods are time consuming and costly. For example Olufunmilayo et al in 2010 evaluated the antisickling activities of *Plumbago zeylanica* and *Uvaria chamae*. Wambebe et al. (2001) reported the use of combination of *Piper guinensis*, *Pterocarpa osun*, *Eugenia caryophylla* and *Sorghum bicolor* extracts for the treatment of SCD. Extracts of *Pterocarpus santolinoides* and *Aloe vera* were reported to increase the gelling time of sickle cell blood and inhibit sickling in vitro (Ugbor, 2006). Sofowora and Isaac- Sodeye (1971) reported the reversal of sickling by root extracts of *Fagara zanthoxylloides*. Thomas et al in 1987 reported the use of an aqueous extract of the unripe fruit of pawpaw (*Carica papaya*) as a treatment for sickle cell disease patients during crises. The research reported the reversal and inhibition of sickling of HbSS red blood cells using aqueous extract. Moody et al. (2003) reported that the aqueous extracts of the reddish brown freshly fallen leaves of *Terminalia catappa* were able to exhibit antisickling activity on sodium metabisulfite induced sickling. Fractionation of an aqueous extract of root bark from *Fagara xanthoxyloides* by column chromatography on DEAE-A-50, utilizing an elution gradient of pH 7.5-5.0, yielded five fractions. All fractions reversed metabisulfite-induced sickling in vitro of erythrocytes homozygous for hemoglobin S (Abu S et al, 1981). Boiled and crude ethanolic extracts of edible *Cajanus cajan* beans were prepared and used for *in vitro* studies involving blood samples obtained from confirmed sickle cell (HbSS) patients. It was demonstrated that the extracts were able not only to inhibit sickling in sodium metabisulfite solution but also quickly reverted to normal morphology of already sickled erythrocytes (Osuagwu CG, 2010).

While all of the aforementioned research investigations were based on prior knowledge of the ethno-medicinal use of the studied plants, in the present investigation we have employed a protocol that relies exclusively on the calculated thermodynamic and structure interaction of over 2000 African natural products with 3D X-ray crystallographic model of human sickle hemoglobin. Because our computer aided drug design (CADD)-guided search for African plants with anti-sickling properties is not based on knowledge of the plants’ ethno-medicinal value, we have been able to conduct a more objective search for the desired activity initialized by virtually screening thousands of natural products before performing antisickling assays. We additionally present an ADME model of the top-performing natural products to properly conceptualize the cell-free and cell-based validation assays presented here.

## Materials and Methods

### 3D natural products library

A number of scientific article databases and repositories including the National Center for Biotechnology Information (NCBI), Pubmed, Chemspider, Pubchem, CHEBI, were searched using different combinations of search terms like “Nigerian plants”, “African plants, “Chemical constituents”, “Chemical structure”, “X-ray”, “Mass spectrometry” and “ Nuclear magnetic resonance”. Articles and records satisfying the search criteria and containing structural information about African natural products were downloaded and further processed. From the 2D structural information of about 2000 natural products covering some two hundred different plant species the 3D structural coordinates were generated using the ligand modeling software ChemBioOffice (Cousins, K.R, 2005). Special attention was accorded to stereogenic sites and the resulting models were subjected to geometric optimization via energy minimization scheme using the integrated MOPAC algorithm (Cousins, K.R, 2005). This helped eliminate steric clashes and allowed each configuration to attain a potential energy minimum. The optimized 3D models were then saved in the Protein Data Bank (PDB) format and pooled together to form the virtual screening library employed in this work.

### Hemoglobin S structure and conformational sampling

The X-ray crystallographic structure (2HBS.pdb) (Harrington et al, 1997) of the dimerized human sickle haemoglobin was downloaded from the RCSB website (https://www.rcsb.org). The structure is a homotetramer involving one tetramer unit (two α and two β globin subunits) interacting with another tetramer unit (Harrington, et al, 1997). The two tetramer units form contact via an interacting surface involving amino acids contributed from two β globin subunits (one from each tetramer). The β globin subunit from Tetramer 1 bears the Glu6Val mutation, while the β globin subunit of Tetramer 2 provides an hydrophobic cavity (both shape and charge complementarity) that Val6 perfectly fits into. The β globin subunit of Tetramer 2 that provides the cavity was thus taken as the receptor molecule since the aim was to competitively inhibit Val6 docking at this site. Preliminary explicit solvent molecular dynamics (MD) simulation had been performed to establish the structural integrity of the Val6 acceptor chain, and the receptor was shown to be stable over a 100 ns MD trajectory performed at 300 K (Olubiyi et al, 2018).

### Virtual screening

Using the software Autodock tool (O.Trott, 2010) and Autodock Vina (O.Trott, 2010), each energy-minimized natural product in our virtual library was docked in a virtual screening protocol into the β globin’s Val6 binding site defined by a hyper-rectangular grid with x, y, z dimension 15.00 Å, 18.75 Å, 15.00 Å. The grid was centered at 6.215, 57.373, 22.171Å to overlap Leu28, Leu42, Leu92, Leu106, Leu110, Leu137, and Leu141, Phe41, Phe71, Phe91, His63, Lys66, and Val67 residues on the Val6 acceptor beta globin molecule and chain B in the crystallographic structure (O.Trott, 2010). The computed binding energies were collected and analyzed to identify the natural products (and hence plants) with the best potential to form thermodynamically stable binding interaction with the HbS lateral aggregation surface.

### Plant collection, botanical authentication and processing

Five out of the seventeen plants identified *via* virtual screening were collected based on availability and within the time frame from different parts of Nigeria. *Rauvolfia vomitoria* Afzel. and *Lantana camara* L. leaves were collected in Agulu, Anambra State Nigeria while the leaves of *Catharanthus roseus* (L.) G.Don, *Hoslundia opposita* Vahl and *Euphorbia hirta* L. were collected in Iyamho, Edo State, Nigeria. They were identified and authenticated at the Department of Botany, Faculty of Sciences, University of Ibadan, Nigeria with voucher numbers 22924, 22925, 22926, 22927 and 22928 for *Hoslundia opposita, Euphrobia hirta, Rauvolfia vomitoria, Lantana camara and Catharanthus roseus* respectively. The plants were deposited in the herbarium of the department. The leaves of the collected plants were air-dried in the laboratory and pulverized into powder using an electrical grinder (750 Watts Euro Premium, Tango DX). 100g of each plant sample was first extracted with n-hexane for three days. Following filtering the residue was extracted with methanol (Analytical grade, Sigma-Aldrich) for 5 days which was then filtered and the filtrate concentrated using a rotary evaporator (DVB RE 100-pro). The concentrates were freeze-dried using a freeze dryer (Zhengzhou Laboao Instrument, LFD-10) and stored at −20°C. The percentage yields were calculated as 7.1%, 1.6%, 5.0%, 5.4% and 1.99% (w/w) for *Catharanthus roseus, Rauvolfia vomitoria, Hoslundia opposita, Lantana camara and Euphorbia hirta* respectively.

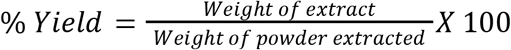

### Antisickling assay

#### Collection of blood

Human blood sample was collected from the residual blood samples obtained from sickle cell clinic at the University College Hospital, Ibadan. The collected blood was centrifuged, washed with phosphate buffer saline pH: 7.4 (PBS: 1.3M NaCl, 0.07MNa_2_HPO_4_ and 0.03M NaH_2_PO_4_) at 4000 rpm for 5 minutes using cold centrifuge.

#### Polymerization Inhibition Test

Polymerization inhibition test was carried out using a modified method previously described by Iwu et al. (1988) and Chikezie et al 2011). This procedure is based on the principle that HbS molecules undergo gelation (gel formation) under deoxygenation conditions induced by the presence of sodium metabisulfite acting as reductant; under this condition deoxyHbS molecules predominate. The degree of polymerization was measured by recording the time-dependent increments in the absorbance of the assay mixture. 0.1 ml of washed HbS was measured into a test tube and equal volume of distilled water was added to lyse the blood cells. 0.5 ml and 1.0 ml volumes of Phosphate buffer saline (PBS) were added to a test tube. The content was transferred to a cuvette and 3.4 ml of 2 % aqueous solution of sodium metabisulfite was added. The absorbance of the mixture was recorded at time zero using a UV/Visible Spectrophotometer (Biobase, BK-UV1800PC) (wavelength = 700nm) and at a 30 seconds interval for a total of 180 seconds (control assay). This procedure was repeated by substituting 1.0 ml of PBS with 1.0 ml of each of the plant extracts at 2 mg/ml, 5 mg/ml and 10 mg/ml) (test assay). Percent polymerization was then calculated as:

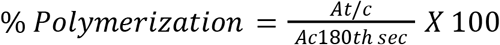

Where: At/c =Absorbance of test/control assay at time = t (s).

Ac180^th^sec = Absorbance of control assay at the 180^th^sec.

#### Sickling reversal test

Sickling reversal test was carried out using the method previously described by Pauline N *et al*, 2013. 20 μL of washed red blood cells was pipetted into a clean Eppendorf tube using a micro pipette. 20 μL of PBS was then added to the Eppendorf tube containing the washed blood. 680 μL of freshly prepared 2% sodium metabisulfite was added and incubated in a thermostated water bath at 37 °C for 1 h. 200 μL of distilled water was added after the initial incubation period and incubated for another 1 h at 37 °C. 10 μL of the incubated cells was transferred onto a hemocytometer. The cells were counted at five different zones (Control assay). This procedure was repeated by replacing 200 μL of distilled water added after the first incubation with 200 μL each of the extract at three different concentrations (10, 5 and 2 mg/ml). The cells were categorized as normal or sickle using visual inspection of their shapes. Biconcave or disk-like shapes were taken to be normal while the elongated, star-like, or wrinkled shapes were considered sickle. The percentage sickle cells were calculated using the following formula:

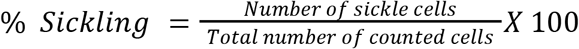

## Results and discussion

### Chemical diversity of the natural products generated

Structures for the 2000 compounds comprising alkaloids, flavonoids, anthraquinones, terpenoids, steroids and coumarins were generated from 200 African plant species (Fig 1). The best performing twenty-three compounds with binding energies ranging from −7.0 kcal to −7.6kcal/mol from seventeen plant species are presented in Table 1

**Figure 1:**
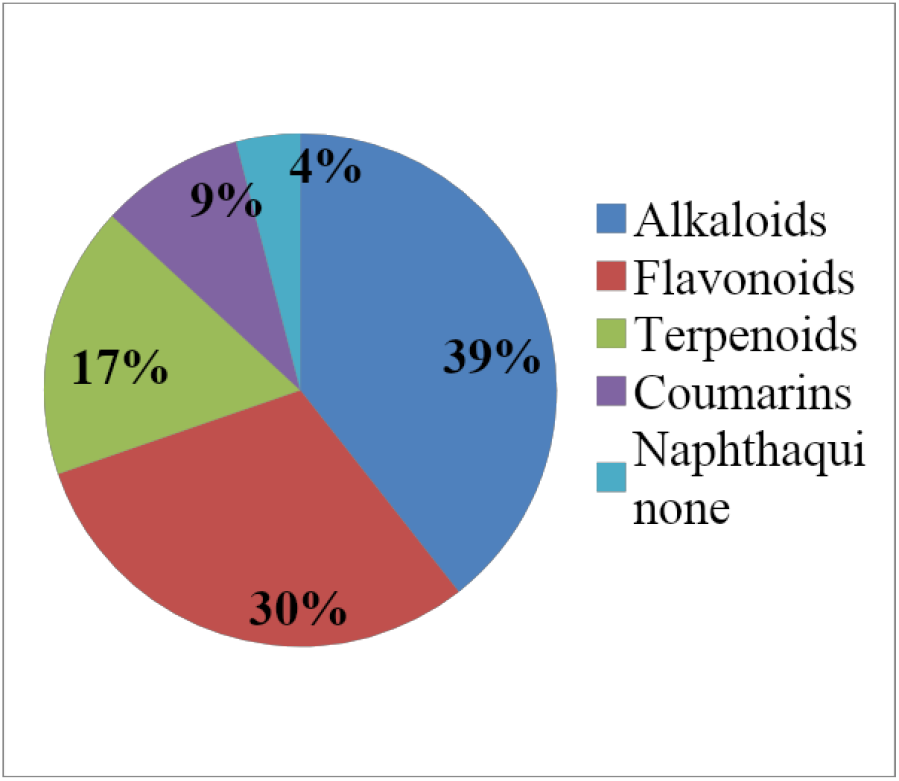
Distribution of the identified compounds into the class of natural products they belong

**Table 1:**
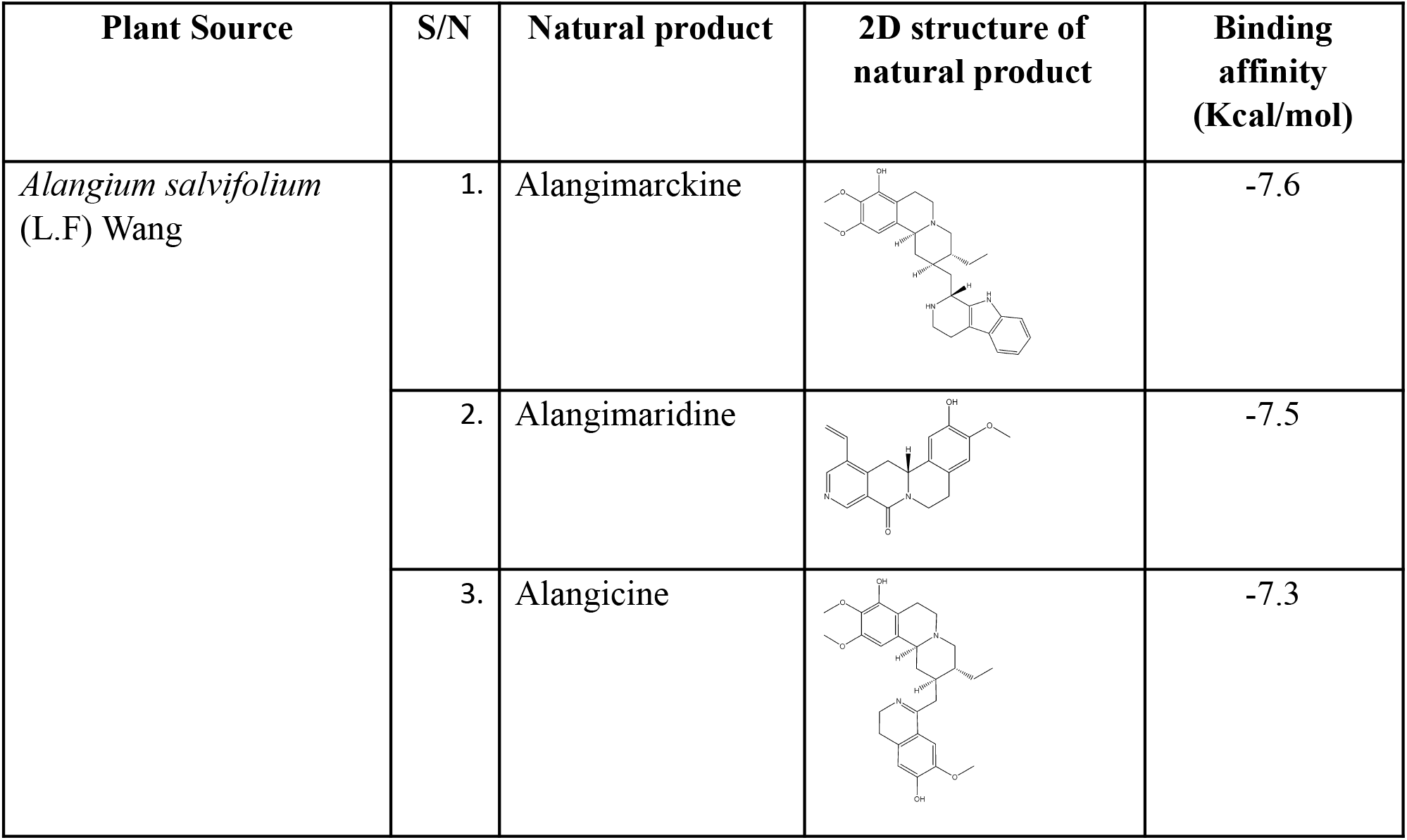
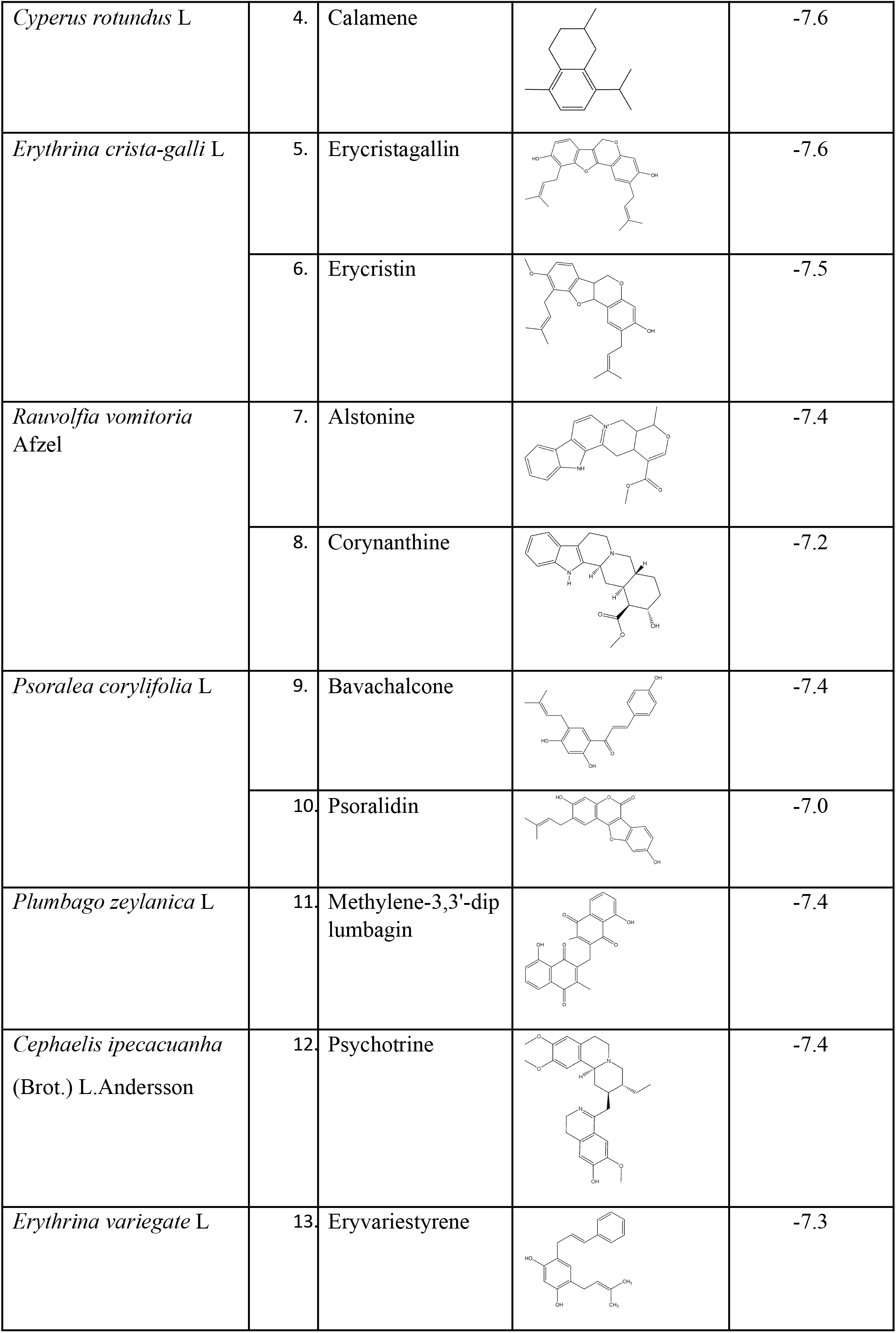
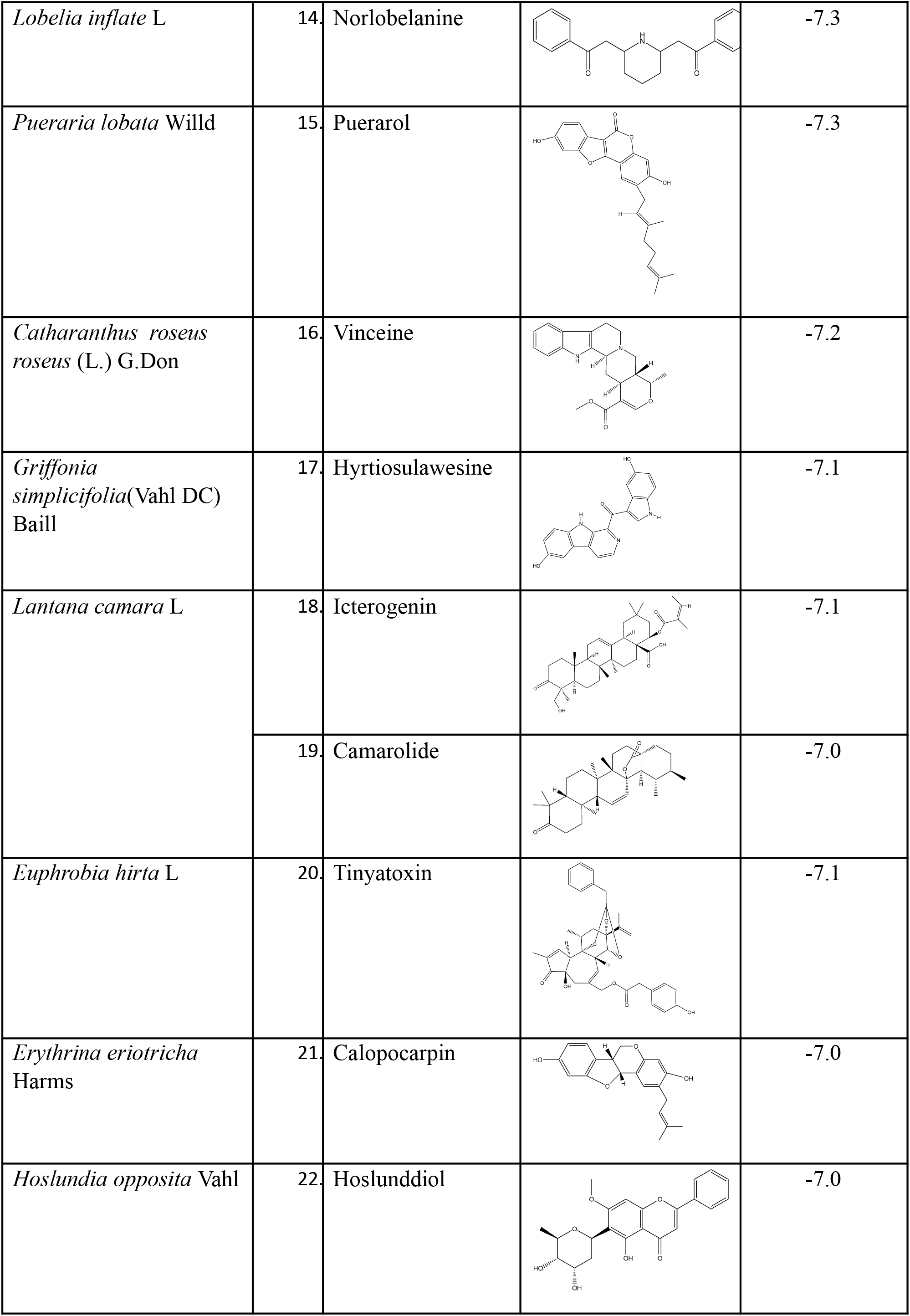
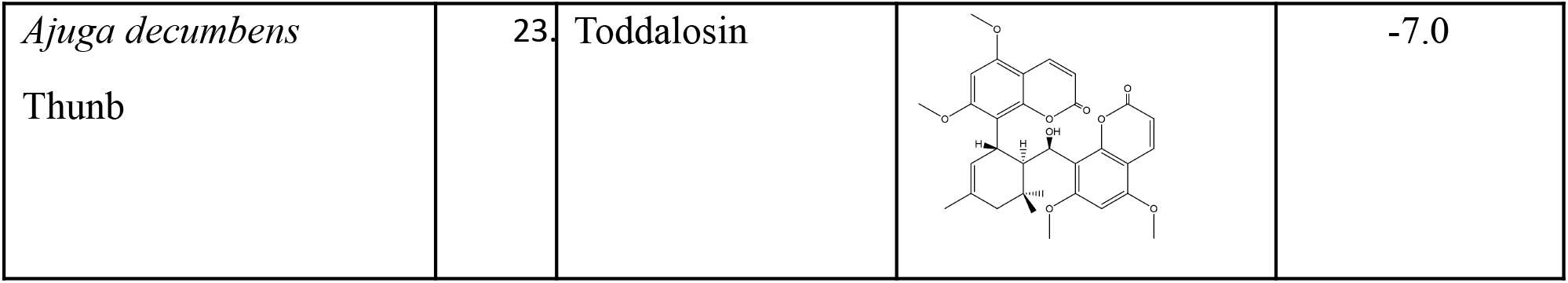
Identified natural products from docking.

It is worthy of note to report that some of the natural products with strong binding HbS interaction identified via *blind* virtual screening in this work are found in plants that had either previously been employed *ethnomedicinally* in SCD treatment or that have been reported to possess antisickling properties. Interestingly, the knowledge of their use in SCD treatment and their anti-sickling properties was unknown to us at the point of natural product library design since the choice of natural products was not based on antisickling properties or use of the plants. For example Olufunmilayo et al in 2010 published a paper in which *Plumbago zeylanica* was confirmed to possess anti-sickling properties. The work however did not offer information about the plant constituent(s) responsible for the observed anti-sickling activity. In the present work, we propose that methylene-3,3’-diplumbagin, a naphthaquinone found in the root of *P. zeylanica* with binding free energy of −7.4 Kcal/mol is possibly one of the constituents responsible for the reported activity. In another work, Egunyomi *et al* (2009) investigated the anti-sickling activities of two ethnomedicinal plant recipes used for the management of sickle cell anemia in Ibadan, Nigeria (Egunyomi et al, 2009) and *Rauvolfia vomitoria* was one of the plants present in the investigated recipes. Two alkaloids, alstonine and corynanthine found in the leaves of the plant exhibited strong computed binding interaction with HbS molecule with binding free energy values of −7.4 and −7.2 kcal/mol, respectively. Their strong binding interaction suggests that they are likely responsible for the earlier reported antisickling activity of the plant. Diosmin and fagarol with binding free energy values of −6.3 and −6.1 Kcal/mol, respectively found in the current exploratory investigation are constituents of *Fagara zanthoxyloides* that had previously been reported by Sofowora *et al*, in 1971 to possess antisickling activity (Sofowora, et al, 1971). Chikezie, reported *Psidium guajava* as possessing HbS polymerization inhibitory at different investigated concentrations. Tamarixetin, gossypetin and guajaverin with binding free energy values of −6.1kcal/mol, −5.8kcal/mol and −5.7 kcal/mol respectively found in the current investigation are constituents of the plant likely to be responsible for the inhibitory effect. Iwu et al (1988) and Ismaila O. Nurain, et al, 2017 in separate investigations reported *Cajanus cajan* as having antisickling activity (74). In our work, two of its constituents, cajanine and cajanulactone were found with binding free energy values −5.9 Kcal/mol and −5.8 Kcal/mol, respectively. We also identified carpaine with binding free energy of −5.9 kcal/mol which is one of the antisickling constituents of *Carica papaya* earlier reported by Thomas *et al*, 1987 and Imaga *et al*, 2009.

### Polymerization inhibition test

The antisickling assay confirmed the ability of the plants collected to both inhibit HbS polymerization and to reduce the extent of RBC sickling. This procedure used to test the antisickling properties of the plant extracts is based on the principle that HbS molecules undergo gelation when deprived of oxygen, transiting to deoxyHbS molecules. Sodium metabisulfite induces polymerization and was hence used as the reductant. The level of polymerization was measured by recording increasing absorbance of the assay mixture with progression of time.

A close look at Table 1 shows that all tested extracts show a reduction in polymerization when compared with the control assay. This displays their anti-sickling properties. Methanolic extract of *Hoslundia opposita* (Table 1) shows a significant (p<0.01) dose-dependent reduction of polymerization to 46.07 ± 0.45%, 74.87 ± 1.20%, and 78.98 ± 0.99% at three different concentrations of 10ppm, 5ppm and 2ppm, respectively. They have *p* values of *p*<0.01 when compared with control. This pattern of polymerization reduction is illustrated in Figure 1. Methanolic extract of *Catharanthus roseus* (Table 2) shows a significant reduction (p<0.01) in polymerization when applied to sodium metabisulfite induced polymerization of HbSS genotype erythrocytes. The reduction was to 75.02 ± 0.54%, 85.82 ±0.35% and 86.32 ± 0.15% at concentrations 10 ppm, 5 ppm and 2 ppm respectively.

**Table 2:** Percentage polymerization of erythrocyte hemolysate of HbSS genotype in the presence of methanolic extracts of *H. opposita, C. roseus and R.vomitoria* at different concentrations.

| Time (s) | Polymerization (%) |  |  |  |  |  |
| --- | --- | --- | --- | --- | --- | --- |
|  | 30 | 60 | 90 | 120 | 150 | 180 |
| Control | 100.00 ± 0.00 | 100.00 ± 0.00 | 100.00 ± 0.00 | 100.00 ± 0.00 | 100.00 ± 0.00 | 100.00 ± 0.00 |
| ***H. opposita*** |  |  |  |  |  |  |
| 10ppm | 45.29 ± 7.78* | 45.90 ± 7.47* | 46.37 ± 7.40* | 46.37 ± 7.54* | 46.50 ± 7.70* | 46.00 ± 7.64* |
| 5ppm | 72.96 ± 3.83* | 74.09 ± 3.20* | 74.69 ± 3.02* | 76.00 ± 2.27* | 75.95 ± 2.45* | 75.53 ± 2.31* |
| 2ppm | 78.15 ± 5.22* | 78.54 ± 5.92* | 78.77 ± 6.13* | 78.69 ± 5.54* | 78.76 ± 5.19* | 80.94 ± 8.58* |
| ***C. roseus*** |  |  |  |  |  |  |
| 10ppm | 74.06 ± 4.15* | 75.49 ± 4.10* | 74.77 ± 6.18* | 75.42 ± 5.52* | 75.30 ± 4.33* | 75.05 ± 5.81* |
| 5ppm | 85.23 ± 4.56* | 85.71 ± 4.48* | 86.22 ± 4.54* | 86.06 ± 4.47* | 85.72 ± 4.56* | 85.97 ± 4.41* |
| 2ppm | 86.45 ± 5.05* | 86.15 ± 4.59* | 86.19 ± 5.26* | 86.23 ± 4.99* | 86.37 ± 4.22* | 86.53 ± 4.11* |
| ***R. vomitoria*** |  |  |  |  |  |  |
| 10ppm | 55.61 ± 15.85* | 54.51 ± 14.55* | 54.26 ± 14.76* | 54.13 ± 14.37* | 53.66 ± 14.26* | 54.17 ± 15.88* |
| 5ppm | 78.64 ± 6.69* | 78.68 ± 6.83* | 78.40 ± 5.90* | 78.43 ± 5.96* | 77.95 ± 6.09* | 77.71 ± 6.63* |
| 2ppm | 83.03 ± 4.10* | 84.38 ± 4.07* | 84.91 ± 4.04* | 84.89 ± 3.95* | 84.53 ± 4.22* | 84.21 ± 4.56* |

Values are expressed as Mean ± Standard deviation of mean. *n*= 5.
\**p*<0.01 (When compared with control).

There was also a significant (p<0.01) reduction of polymerization of erythrocyte hemolysate of HbSS genotype in the presence of methanolic extract of *Rauvolfia vomitoria vomitoria* at different concentrations as presented in table 2. It reduces polymerization to 54.39 ± 0.66%, 78.30 ± 0.39% and 84.49 ± 0.36% for 10ppm, 5ppm and 2ppm respectively. In all there was an overall significant reduction of polymerization by the extracts of the different plant species tested. This goes to show that the plants have the ability to inhibit polymerization at different concentrations of 10ppm, 5ppm and 2ppm.

We obtained good dose-dependence for the polymerization assay in which the natural products in the different extracts have unfettered and direct access to the hemoglobin molecule and to the aggregation sites (Figure 3). In the sickling test however, this is not so: there is a layer of complexity, the natural barrier introduced by the plasma membrane (Table 3). This barrier constituted by the cell membrane in the sickling test most likely poses a pharmacokinetic challenge to the natural products, and prevents quite a number of their molecules from crossing into the RBC core where the hemoglobin molecules are located. Thus, the observed activity in the antisickling test (being cell based) is extensively modified by cell membrane penetration.

**Figure 2:**
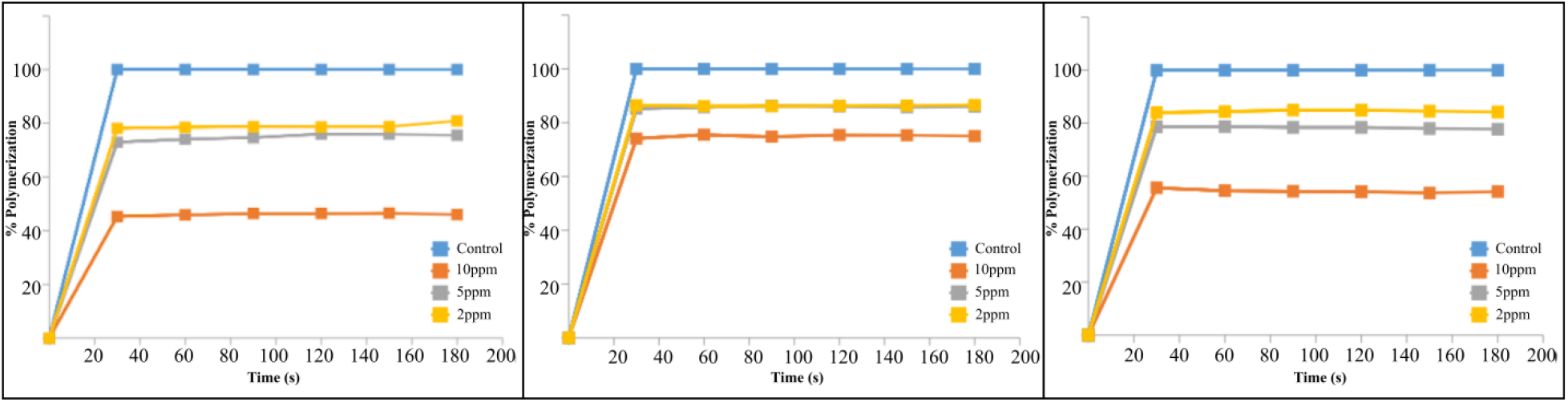
Percentage polymerization of erythrocyte hemolysate of HbSS genotype in the presence of methanolic extract of *Hoslundia opposita, Catharanthus roseus and Rauvolfia vomitoria* at three different concentrations. Values are expressed as Mean ± Standard deviation of mean. *n*= 5. ^*^p<0.01 (When compared with control).

**Figure 3:**
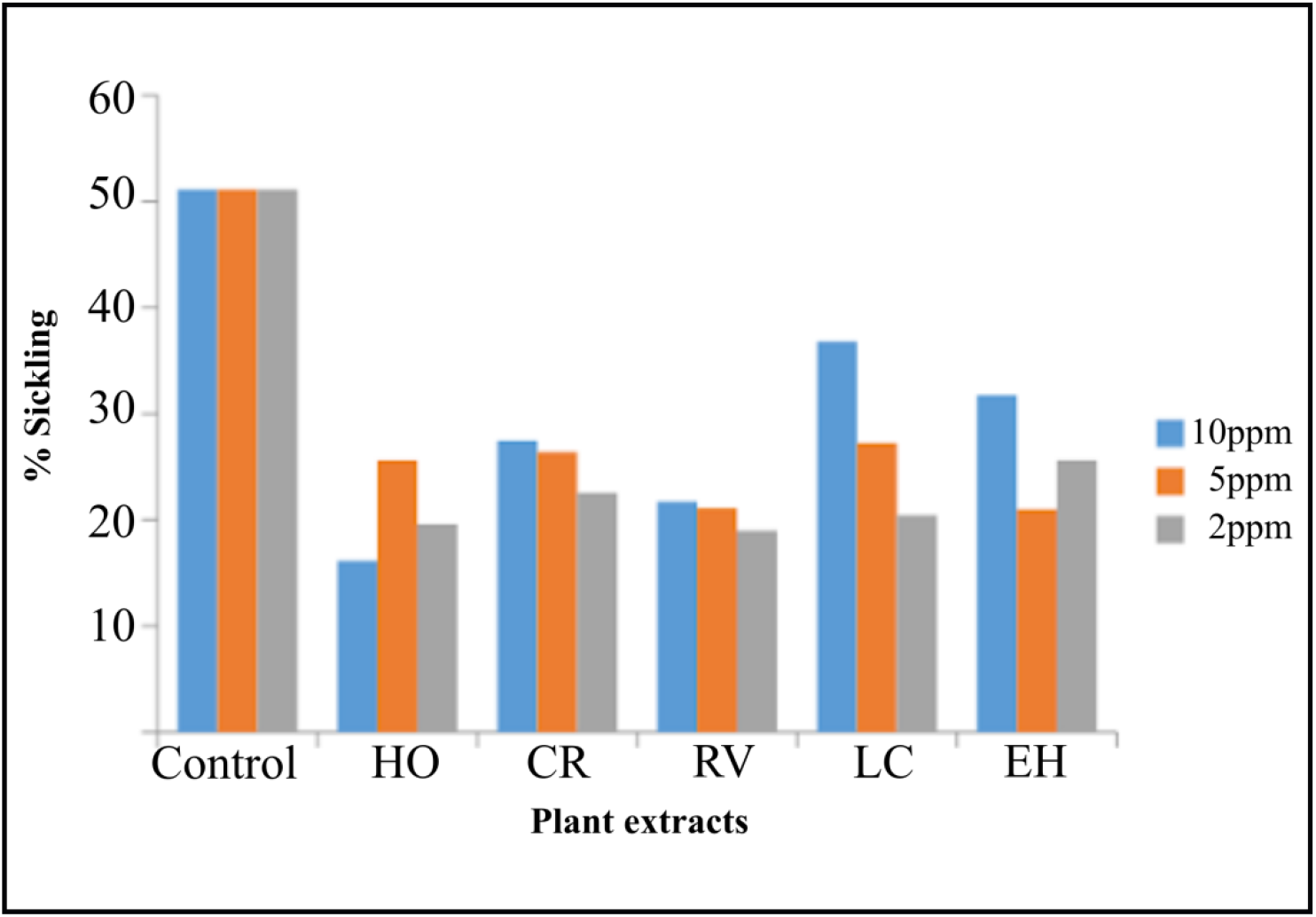
Percent Sickling of RBC in the presence of different plant extracts.

**Table 3:** Percentage sickling of RBC in the presence of different concentrations of the five different extracts.

| Sample | % Sickling of RBC |
| --- | --- |

|  | **10ppm** | **5ppm** | **2ppm** |
| --- | --- | --- | --- |
| Control | 51.10 ± 6.80 | 51.10 ± 6.80 | 51.10 ± 6.80 |
| *Hoslundia opposita* | 16.12 ± 2.10* | 25.61 ± 3.25* | 19.60 ± 2.54* |
| *Catharanthus roseus* | 27.47 ± 6.17* | 26.39 ± 5.78* | 22.57 ± 2.19* |
| *Rauvolfia vomitoria* | 21.70 ± 4.40* | 21.12 ± 4.02* | 18.98 ± 6.10* |
| *Lantana camara* | 36.79 ± 5.72* | 27.21 ± 3.76* | 20.42 ± 2.53* |
| *Euphorbia hirta* | 31.74 ± 3.74* | 20.99 ± 3.58* | 25.61 ± 9.27* |

Values are expressed as Mean ± Standard deviation of mean. *n*= 5.
\**p*<0.01 (When compared with control).

We believe that their polymerization inhibition capacity can be attributed to the phytochemicals earlier identified to be present in the various plant species and docked at the hydrophobic pocket of the HbS genotype. *Hoslundia opposita* contains the phytochemicals hoslunddiol, a flavonoids having binding energy of −7.0kcal/mol. A close look at the structure of hoslunddiol reveals the presence of phenyl (C_6_H_5_) group that helps to bind strongly at the hydrophobic pocket of the HbS genotype hence blocking the binding of valine to it. It also possesses a hydrophilic hydroxyl side chain that helps to disrupt the docking of Val6 on an incoming beta globin chain. This characteristic possessed by hoslunddiol is likely to be responsible for the antisickling activity displayed by the extract of *Hoslundia opposita*.

Similarly, *Catharanthus roseus* contains the phytochemical vinceine (ajmalicine), an alkaloid found in the leaves of the plant and having a binding energy of −7.2kcal/mol. The structure of vinceine has a non-polar benzene (C_6_H_6_) ring for binding and an acetate (CH_3_COO^−^) hydrophilic side chain helping to disrupt the docking of Val6 on an incoming beta globin chain. This is what is likely to be responsible for the antisickling activity shown by *Catharanthus roseus*.

*Rauvolfia vomitoria* has two phytochemicals of interest, astonine and corynanthine, both alkaloids with binding energy −7.4kcal/mol and −7.2kcal/mol respectively. Their structures both show the presence of a non-polar benzene (C_6_H_6_) ring for binding and an acetate (CH_3_COO^−^) hydrophilic side chain helping to disrupt the docking of Val6 on an incoming beta globin chain. This study reveals that, even though Alstonine has the lowest binding energy of −7.4kcal/mol compare to hoslunddiol with binding energy −7.0 Kcal/mol, *Hoslundia opposita* at 10ppm has a greater percentage (53.93%) of polymerization inhibition compare to 45.61% and 25.46% polymerization inhibition of *Rauvolfia vomitoria* and *catharanthus roseus*. This could be likely due to the presence of more electronegative functional groups in alstonine thereby making it to have a more hydrophilic side chain that helps to disrupt the docking of Val6 on an incoming beta globin chain (Figure 4). Refer to table 1 for the structures of the phytochemicals of interest.

**Figure 4:**
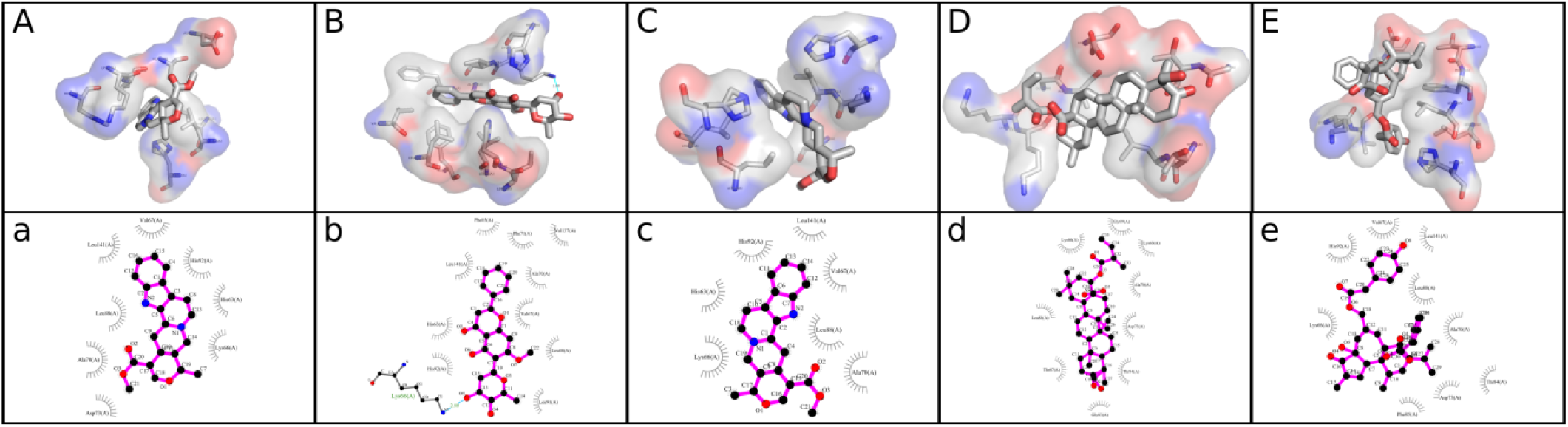
Interaction between the best compound from each of the five plants screened and the HbS receptor. The order of arrangement from left to right is: Alstonine, Hoslunddiol, Vinceine, Icterogenin and Tinyatoxin.

All five tested extracts show significant (p<0.01) reduction of sickling of red blood cells (RBC) after inducing sickling with sodium metabisulfite. Methanolic extract of *Hoslundia opposita* shows a significant reduction of sickling after inducing sickling with sodium metabisulfite by 16.12 ± 2.10, 25.61 ± 3.25 and 19.60 ± 2.54 at concentrations 10ppm, 5ppm and 2ppm respectively when compare with control having a 51.10% sickling. Methanolic extracts of *Catharanthus roseus* and *Rauvolfia vomitoria* also show a significant reduction in sickling by 27.47 ± 6.17, 26.39 ± 5.78, 22.57 ± 2.19 for *C. roseus* and 21.70 ± 4.40, 21.12 ± 4.02, 18.98 ± 6.10 for *Rauvolfia vomitoria* at concentrations 10ppm, 5ppm and 2ppm respectively. Similarly, chloroform extract of *Lantana camara* L reduces percentage sickling of sickle red blood cells by 36.79 ± 5.72, 27.21 ± 3.76 and 20.42 ± 2.53 at concentrations 10ppm, 5ppm and 2ppm respectively. Chloroform extract of *Euphorbia hirta* also reduces percentage sickle of red blood cells when induced with sodium metabisulfite by 31.74 ± 3.74, 20.99 ± 3.58 and 25.61 ± 9.27 at concentrations 10ppm, 5ppm and 2ppm respectively.

*Hoslundia opposita* at 10ppm, 5ppm and 2ppm, reverse sickling of sickle red blood cells by 68.50%, 49.89% and 61.64% respectively *Catharanthus roseus* reverses sickling of sickle red blood cells by 46.24%, 48.35% and 55.84% at concentrations 10ppm, 5ppm and 2ppm respectively. Similarly, we observed that extract of *Rauvolfia vomitoria* shows a reversal in sickling of RBC by 57.53%, 58.67% and 62.85% for the three different working concentrations. There was also a reversal of sickling of sickle red blood cells by extracts of *Lantana camara* and *Euphorbia hirta* by 28.00%, 46.75% and 60.04% for *L. camara* and 37.88%, 58.92% and 49.88% for *E. hirta* at concentrations 10ppm, 5ppm and 2ppm respectively.

It is observed that the extracts show different capacity of sickling reversal at the different concentrations. For example, extract of *H. opposita* shows a greater percent (68.50%) of sickling reversal at 10 ppm and the least percent (49.89%) at 5ppm. *Catharanthus Roseus* (L.) G.Don displayed a greater percent (55.84%) of sickling reversal at 2ppm and shows the least percent (46.24%) at 10ppm. The highest percent sickling reversal for *Rauvolfia vomitoria* Afzel was observed at 2 ppm (62.85%) while the smallest was observed at 10 ppm (57.53%). Similarly, *Lantana camara* L displayed the least percent (28.00%) of sickling reversal at 10ppm and the most percent (60.04%) at 2ppm while extract of *Euphorbia hirta* L has the least percent (37.88%) of sickling reversal at 10ppm and the most percent (58.92%) at 5ppm (Table 3).

A close look at Table 3 and Figure 3 shows that at 10ppm, extract of *Hoslundia opposita* Vahl has the highest percent of sickling reversal (68.50%) when compared to others while extract of *Lantana camara* L shows the least percent sickling reversal (28.00%). At 5ppm extract of *Euphorbia hirta* L shows the most percent (58.92%) sickling reversal of sickle red blood cells while extract of *Lantana camara* L shows the least percent (46.75%) sickling reversal. Similarly, extract of *Rauvolfia vomitoria* Afzel shows the most percent (62.85%) sickling reversal of RBC at 2ppm while extract of *Euphorbia hirta* L shows the least percent (49.88%) reversal of sickling of sickle red blood cells. The overall most significant reduction of sickling for the different extracts and concentration was observed for *Hoslundia opposita* Vahl (68.50%) at 10ppm while the overall least percent (28.00%) sickling reversal was observed for extract of *Lantana camara* L at 10ppm.

Lastly, we compared surface maps for the static molecular features of Glu and Val (Figure 5) which reveal subtle differences between both *mutants* that we believe underlie the distinctions in the aggregation behaviour of HbA and HbS. From the presented electronic distribution, electrostatic and lipophilicity maps, the aggregation observed in HbS (and thus RBC sickling) is likely a consequence of the differences observed in the electrostatic and lipophilic features of Glu and Val. The pathologic Glu6Val mutation replaces an electron-rich glutamate side-chain capable of electrically preventing hydrophobic aggregation to a highly lipophilic valine side-chain that encourages aggregation. It is interesting to note that such seemingly minor shape and charge perturbation resulting from the point mutation is capable of provoking such far reaching structural and aggregational consequences that underlies SCD symptom complex. Perhaps the seemingly disproportionate conformational consequence of the mutation partly results from the allosteric nature of the hemologbin molecule which allows tiny changes to be amplified in processes downstream of the point of mutation. It is difficult to categorically state whether such influence also plays a part in the inhibitory action of the plant extracts studied in the present work. Analysis of the natural product-HbS interaction indeed reveals a good fit in Figure 4 and we believe that the inhibitory activities observed is at least partly a consequence of the interaction between the natural products and the investigated binding site of the sickle hemoglobin molecule. A categorical attribution of the inhibitory activities will however require a more detailed structural characterization of the binding event using spectroscopic characterization, especially X-ray spectroscopy. This is presently beyond the scope of this work whose aim is to present a procedure for employing natural products-based virtual screening in identifying plants with anti-sickling activities.

**Figure 5:**
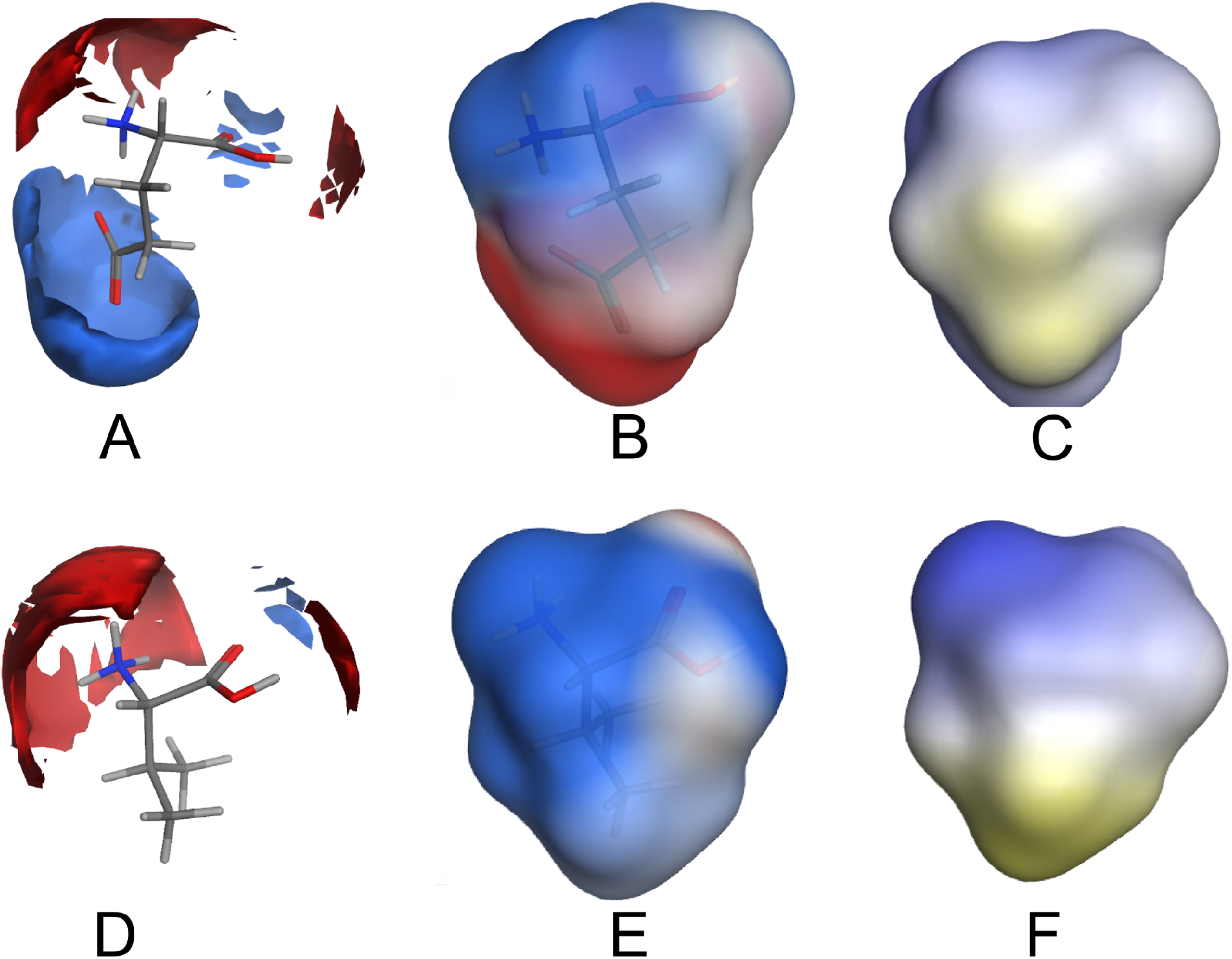
Surface maps comparing electronic charge distribution around Glu vs Val, electrostatic distribution lipophilicity surface maps of Glu vs Val. A.) Electronic charge interaction distribution around Glu showing positive charge preference around the carboxylate group. B.) Electrostatic potential distribution on Glu. C.) Lipophilicity surface map of Glu. D.) Electronic charge interaction distribution around Val. E.) Electrostatic potential distribution on Val. C.) Lipophilicity surface map of Val. Blue, red and light colors in electronic charge interaction and electrostatic potential maps indicate positive, negative and neutral charges, respectively; while light to yellowish color indicate increasing lipophilic nature in the lipophilic surface maps. The computed LogP(o/w) values for zwitterionic glutamate and valine are −0.844 and 0.313, respectively.

### Pharmacokinetics considerations

It is important to consider that effects obtained in the cell-based antisickling experiment here likely has the involvement of pharmacokinetic contributions especially with respect to gaining access to the intracellular target. Focusing on the natural products with the best interaction with HbS we performed ADME modeling using the SwissADME web-based application (Daina and Zoete, 2016; Daina and Zoete, 2017) and focusing on gastrointestinal absorption as an indication of ability to cross cell membranes. Out of the 22 natural products so profiled, only three-calamene, icterogenin and toddalosin- were predicted to be associated with poor gastrointestinal (GIT) absorption. Apart from the three compounds being fairly *strongly* lipophilic (Log P ranging from 4.9 to 7.1), calamene is a hydrocarbon expected to be associated with aqueous solubility liabilities while the other two have molecular weights in excess of 500 g/mol. These features are likely responsible for the predicted defective absorption in the GIT. Transport across erythrocytic plasma membrane involves mobility across membrane bilayer essentially similar to that across the GIT epithelial cells and the predicted GI transport profiles may be taken to imply the compounds potential to cross RBC plasma membrane and gain access to intracellular HbS. Based on this nineteen natural products with high GI transport features are expected to attain intracellular concentrations sufficient for HbS aggregation. However, 50 % of the compounds have the potential to cross the blood brain barrier (BBB) and might be associated with central nervous system effects. Even though it is only speculative what the role of P-glycoprotein (Pgp) binding will be in sickle cell disease drug management, we have included the Pgp interaction profiles for the natural products as their use would have implications in co-morbidities where Pgp inhibition is important. Majority of the natural products were predicted to possess Pgp inhibitory potential including hoslunddiol, icterogenin, vinceine, and tinyatoxin that are respective constituents of *H. opposita*, *L. camara*, *C. roseus*, and *E. hirta* tested in this work. We expect that this would have implications for the use of these natural products and the corresponding plants in SCD management involving (for example) cardiovascular comorbidities. It is interesting to note that alstonine (in *R. vomitoria*) was predicted to be unlikely to inhibit Pgp. We similarly obtained the predicted interaction profiles for five key metabolic enzymes (CYP1A2, CYP2C19, CYP2C9, CYP2D6, and CYP3A4) likely to influence their metabolic disposition as well as interaction with co-administered drugs or herbal products. Out of the five natural products found in the plants that we tested only the compound icterogenin (*L. camara*) shows zero likelihood of inhibiting any of the five cytochrome P450 enzymes. Hoslunddial (*H. oposita*) shows potential for inhibiting CYP3A4; vinceine (*C. roseus*) will likely inhibit CYP2D6, while alstonine (*R. vomitoria*) shows possibility of inhibiting both enzymes as well as CYP2C19. Tinyatoxin was predicted with CYP2C9 and CYP3A4 inhibitory activity but none of the compounds was shown with predicted inhibitory action on CYP1A2. It however remains to be examined how these profiles will translate to clinical significance and if the titres of these natural products administered (e.g. in herbal preparations) constitute strong ground for meticulous pharmacotherapy review.

## Conclusion

We have presented a protocol that employs virtual screening in identifying natural products with sufficiently strong interaction strength with the sickle hemoglobin. This guided our collection of plant parts and subsequent experimental testing of hemoglobin polymerization inhibitory activities as well as antisickling properties in whole cells. We have thus shown that, beyond the limitations imposed on ethnomedicinal research as a result of dependence on only ethnic knowledge and experience, our approach is capable, through screening of several natural products, of identifying plants with anti-sickling properties. It also has the advantage of achieving environmentally friendly lead finding research which assures the collection of only the plants with a good likelihood of producing desired pharmacological activities thereby reducing wastage of plant materials. Our approach has also helped us to gain time through screening of thousands of compounds within a short period. It is the first time deploying CADD for the identification of plants with antisickling activities in sickle cell disease research. The outcome of our work will be of importance both for development of new leads for SCD drug development.

## Authors contributions

**Eric O. Olori**: 3D Library Design, Virtual Screening, Plant Collection and In vitro testing. Writing- Original draft preparation. **Olujide O. Olubiyi**. : Conceptualization, 3D Library Design, Virtual Screening, Data curation, Writing-Original draft preparation, Editing. **Chinedum P. Babalola**: Conceptualization, Project Planning and Supervision, Writing-Original draft preparation, Reviewing and Editing.

